# Dominant suppressor genes of p53-induced apoptosis in *Drosophila melanogaster*

**DOI:** 10.1101/2024.02.06.579196

**Authors:** Tamás Lukacsovich, Tamás Szlanka, Éva Bálint, Kornélia Szabó, Ildikó Hajdu, Enikő Molnár, Erika Virágh, Yu-Hsien Lin, Orsolya Méhi, Ágnes Zvara, István Török, Zoltán Hegedűs, Brigitta Kiss, Beáta Ramasz, Laura M. Magdalena, László Puskás, Bernard M. Mechler, Adrien Fónagy, Zoltán Asztalos, Michal Žurovec, Imre Boros, István Kiss

## Abstract

Apoptosis, the programmed cell death, is responsible for the removal of cells seriously damaged or unwanted in development. A major function of apoptosis is the removal of cells which suffered oncogenic mutations, thereby preventing cancerous transformation.

By making use of the *DEP* transposon, a *P* element derivative made in our laboratory, we made an insertional mutagenesis screen in *Drosophila melanogaster* to identify genes which, when overexpressed, suppress the *p53*-activated apoptosis. The *DEP* element has Gal4-activatable, outward-directed *UAS*-promoters at both ends which can be deleted separately *in vivo*. In the *DEP* insertion mutants, we used the *GMR-Gal4* driver to induce transcription from both *UAS*-promoters and tested the suppression effect on the apoptotic rough eye phenotype generated by an activated *UAS-p53* transgene.

By *DEP* insertions, seven genes were identified which suppressed the *p53*-induced apoptosis. In four mutants, the suppression effect was resulted by single genes activated by one *UAS*-promoter (*Pka-R2, Rga, crol, Spt5*). In the other three (*Orct2, Polr2M, stg*), deleting either *UAS*-promoter eliminated the suppression effect. In qPCR experiments we found that the genes in the vicinity of the *DEP* insertion also showed an elevated expression level. This suggested an additive effect of the nearby genes on suppressing apoptosis.

In the eucaryotic genomes there are co-expressed gene clusters. Three of the *DEP* insertion mutants are included and two are in close vicinity of separate co-expressed gene clusters. This raises the possibility that the activity of some of the genes in these clusters may help the suppression of the apoptotic cell death.

## Introduction

Cells seriously damaged by stress or not needed in development are removed by the process of regulated cell death: apoptosis, autophagy and necroptosis (Galluzzi *et al*. 2015). Apoptosis, plays a critical role in both normal biological processes requiring cell removal (Arya and White 2015).and stress responses causing badly damaged cells to undergo apoptosis.

The central mediator of the stress response is *p53*, a transcription factor, which can induce cell cycle arrest or programmed cell death depending on the extent of damage. The *p53* gene acts as a central point of integration receiving diverse signals of stress and damage, and responding to them by a plethora of changes in gene regulation, cell signaling and direct molecular interactions, resulting in the activation of the cell cycle checkpoints and/or the apoptosis cascade (Vousden and Lane 2007; Levine and Oren 2009; Inoue *et al*. 2016).

A major activator of the *p53* gene is the genetic stress (DNA damage, lethal mutations, aneuploidy, telomere shortening, aberrant gene activation) which could lead to aberrations in the cell cycle and uncontrolled cell proliferation and cancer. By killing abnormal cells through apoptosis, *p53* acts as a „guardian of genome integrity” which prevents tumorous transformation of the cells *(Albrechtsen* et al. *1999; Mehta* et al. *2012)*. In fact, *p53* is one of the most important tumor suppressor genes *(Olivier* et al. *2010; Muller and Vousden 2013)*.

In addition to the genetic pathways implementing apoptosis, numerous downstream genes have been identified which are regulated by *p53*. Less is known about the upstream regulation of *p53* and apoptosis (Shen and White 2001; Menendez *et al*. 2009; Carvajal and Manfredi 2013). With respect to the cancerous transformation, the negative regulators/suppressors of *p53* and/or apoptosis are of particular importance (Portt *et al*. 2011). Under normal conditions, such genes prevent unwanted cell death. However, their abnormally elevated expression may interfere with the normal regulation of apoptosis, opening the gate to abnormal cell proliferation and cancer progression (Hollstein and Hainaut 2010; Olivier *et al*. 2010; Lui *et al*. 2013; Wang *et al*. 2014).

Since the discovery of a *p53* orthologous gene in *Drosophila* (Ollmann *et al*. 2000; Jin *et al*. 2000; Brodsky *et al*. 2004), *Dmp53* has been used as a useful model to study *p53* function and regulation. Although the DNA sequence homology with the mammalian *Tp53* is not particularly high, the protein structure, the function and the interaction network of the mammalian and fruit fly orthologs are evolutionarily conserved and largely similar, therefore the results gained in *Drosophila* can easily be interpreted for the mammalian system (Steller 2008; Rutkowski *et al*. 2010; Papagiannouli *et al*. 2013; Mollereau and Ma 2014).

Large-scale screening of the genomes of model organisms for mutations with special phenotypes is a powerful approach for the analysis of complex biological functions (Cooley *et al*. 1988; Török *et al*. 1993; Rørth 1996). By controlled overexpression of specific genes, one can identify important genetic interactions. The strategy of selectively activating random genes by the insertion of P element constructs which carry Gal4-inducible promoters, e.g., the *EP* element (Rørth 1996) or the *GS* construct (Toba *et al*. 1999), was successfully applied previously. To recover dominant suppressors of *p53*-induced apoptosis, we made a gain-of-function screen in *Drosophila melanogaster* by making use of the *Gal4*-activitable *DEP* P element transposon made in our laboratory. We identified seven insertion mutants which, when overexpressed, significantly suppressed the apoptotic effect in the eyes, the rough eye (*r.e.*) phenotype, in the *GMR-Gal4>UAS-p53* combination. In three of them, however, the activation of one gene was not enough to exert the suppression effect. As the genes around the *DEP* insertion are also activated to some extent, in these cases the suppression of apoptosis is probably resulted by an additive effect including the genes in close vicinity. Regarding the suppression of apoptosis, this type of additive activity had not been observed previously.

In the eucaryotic genomes there are gene clusters in which the neighbor genes are co-expressed. Some of the clusters contain genes with similar functions, while others have genes with diverse functions. We compared the chromosomal location of the genes in the *DEP* insertion neighborhoods with that of the known co-expressed gene clusters (Spellman and Rubin 2002) in the *Drosophila* genome. Interestingly, we found that three of the *DEP* insertion mutants were included in three separate co-expressed clusters, while two mutants were located outside but very near to the boundary of two further clusters.

This paper describes the isolation and preliminary characterization of these apoptosis suppressor genes in *Drosophila melanogaster*.

## Materials and Methods

### Fly cultures and stocks

Fly cultures were kept on standard cornmeal-yeast-agar medium at 25°C if not otherwise stated. The genetic combinations tested were established by standard genetic crosses on *w* homozygous background. The ***P(Δ2-3)***: *ry[506] P{ry[+t7.2]=Delta2-3}99B*, ***GMR-Gal4****: w[*]; P{w[+mC]=GAL4-ninaE.GMR}12, **UAS-p53***: *y[1] w[1118]*; *P{w[+mC]=UAS-p53.H159N.Ex}3*, ***yw; MKRS, FLP/TM6B, Cre***: *y[1] w[67c23]; MKRS, P{ry[+t7.2]=hsFLP}86E/TM6B, P{w[+mC]=Crew}DH2, Tb[1*], ***UAS-stg***: *w[1118]; P{w[+mC]=UAS-stg.N}16/CyO, P{ry[+t7.2]=sevRas1.V12}FK1* and *w[*]; T(2;3)ap[Xa], ap[Xa]/CyO; TM6* stocks were received from the BDSC Stock Center, Bloomington, Indiana. The shortened genotypes in bold preceding the complete ones represent the name used in the text. *UAS-Spt5* was a kind gift from Ruth Palmer. Transgenic *RNAi* stocks were received from the NIG-FLY (Mishima), VDRC (Vienna) and BDSC (Bloomington) collections. The *w, DEP* homozygous stock used for the transposon mutagenesis was created in our laboratory; see below. In the description of the genetic constructs, we followed the terms of Flybase (http://flybase.org).

### Construction of the DEP activating transposon

The *pDEP* construct was made in our laboratory as follows: At first we replaced the entire genetrap cassette in the backbone of *pGT1* vector (Lukacsovich *et al*. 2001) with the *mini-white**^+^*** (*m-w**^+^***) gene of *pCasper2*. This step resulted in unique NotI as well as XhoI restriction sites next to the 5’ and 3’ P-element ends, respectively. Using these sites, two Multi-Cloning Sites (MCS) containing several unique restriction sites were inserted on both sides of the *m-w**^+^*** gene by ligating synthetic double stranded oligonucleotides into the locations. The *5xUAS-hsp70*-core promoter fragment – from the *pUAST* vector, (Brand and Perrimon 1993) – and the *loxP* and *FRT* sequences were then inserted in the desired orientations into the MCSs to get the final *DEP* construct (**Fig. 1**). A detailed description of the steps of construction is available upon request. As Figure1 shows, the sequence unit containing the *UAS* promoter at the 5’ end of *DEP* and the *m-w**^+^*** gene together are flanked by *FRT* sequences (*UAS^FRT^*) while the 3’ *UAS* and the *m-w**^+^*** are between two *loxP* sites (*UAS^loxP^*). This arrangement makes the *UAS* promoters selectively deletable *in vivo* by the FLP or Cre recombinases. The *pDEP* construct was microinjected along with the *Δ2–3* transposase helper plasmid into *w ^1118^* syncytial blastoderm-stage embryos by using standard techniques. Surviving adults were crossed again to *w* homozygous flies, and in the next generation transformants were screened for their red eye color, and X-chromosomal insertions were selected.

### Genetic screen for dominant modifiers of the p53-induced apoptosis

As shown in Figure S1, female flies carrying the *DEP* element on the X chromosome were crossed to males of the *P(Δ2-3)* jumpstarter stock producing the *P* element transposase (Robertson *et al*. 1988). Remobilized by the transposase, the *DEP* element „jumps out” of the X chromosome and gets inserted at new sites in the genome. Males carrying the new insertions in their germline were crossed to females of a *T(2;3)* translocation balancer *w/w; T(2;3)ap[Xa], ap[Xa]/CyO; TM6*. In the next generation, male offspring (*w/Y*) have white eyes, except those which carry new autosomal *DEP* insertions and have colored eyes by the *m-w**^+^***expression. Single males with colored eyes were simultaneously crossed to *T(2;3)* translocation balancer females (see above) and homozygous „tester” females of *w; GMR-Gal4; UAS-p53* genotype. In the next generation, if the *DEP* insertion mutant activated by the *GMR-Gal4* driver suppressed the *p53*-induced „rough-eye” (*r.e.*) phenotype, red-eyed males carrying the new *DEP* suppressor mutant above *CyO* or *TM6* balancer were crossed again to the appropriate balancer females. Through serial crosses to balancer stocks, the new insertions on the 2nd or 3rd chromosomes were isolated as homozygous mutant lines.

### Determination of DEP insertion sites

Inverse PCR was performed according to the protocol described previously (Kyriacou 2000), with some modification. Shortly, genomic DNA of approximately 10 flies was extracted, digested with restriction enzyme HpaII (NEB) and after phenol-chloroform extraction the resulting fragments were ligated with T4 ligase (NEB) for 2 hours at room temperature to circularize them. Two μl out of the 20 μl ligation mixture was used as template in the PCR reaction. The PCR reactions were performed using Taq DNA polymerase (Qiagen) with the Taq PCR buffer, 1.5 mM MgCl2, 0.2 mM dNTPs and a primer pair specific to the 3’P-end of the DEP element: P3’Fw1 that hybridizes between nucleotide positions 106 and 131 in the DEP vector (GTCTGAGTGAGACAGCGATATGATTG) and P3’Rev1 which binds to the vector between positions 75 and 51 (CACTCGCACTTATTGCAAGCATACG) on the complementary strand, both at 0.5 μM final concentration. The sample was cycled 35 times for 30 seconds at 95 °C, 30 seconds at 58 °C and 1 minute at 72 °C. 1μl of the resulted reaction mixture was used as template for a second round of PCR reaction using the following nested primer pair: P3’Fw2 that hybridizes between nucleotide positions 131 and 154 (GTTGATTAACCCTTAGCATGTCCG) and P3’Rev2 which binds to the vector between positions 50 and 28 (TTAAGTGGATGTCTCTTGCCGAC) on the complementary strand, again at 0.5 μM final concentration. The second-round reaction was performed under the same conditions as the first round except the annealing temperature was elevated to 60 °C. After purification (Qiagen QIAquick) the PCR product was sequenced with primers P3’Fw2 and/or P3’Rev2. Sequence data were blasted to FlyBase (*D. melanogaster*, r6.49) to identify the genomic region carrying DEP insertion. Insertion points were verified in a third round of PCR reaction using a primer specific to the 5’P-end of the DEP element (P5’Fw) that hybridizes between nucleotide positions 5483 and 5504 in the DEP vector (GTA TAC TTC GGT AAG CTT CGG C) and a primer specific to each of the relevant genomic regions identified. The *DEP* insertion site sequences are given in the **Supplementary Table 1**.

### Selective in vivo deletion of the UAS promoters in the DEP insertion mutants

To induce promoter deletion in the *DEP* element, the suppressor mutants (*suppr^DEP^*) were crossed to *yw; MKRS, FLP/TM6B, Cre* flies, where the *MKRS* and *TM6B* balancer chromosomes carry heat-inducible transgenes of the FLP and Cre site-specific recombinases, respectively. The recombinases were induced by heat shock (37°C, 2 hrs) in 2nd instar larvae of the F1 generation. The male F1 flies carrying the *MKRS* or *TM6 B* chromosomes were separately crossed to homozygous *w* balancer stocks. Because the *UAS* (along with the coupled promoter) and the *mini-w^+^* marker were removed together, the *UAS*-deleted flies (*suppr^DEPΔFRT^* or *suppr^DEPΔloxP^*) in the next generation could be recognized by the white eye color (**Fig. 1**).

### Silencing the suppressor genes with RNAi

To test whether silencing the *DEP*-bearing gene really weakened the suppression of apoptosis, we constructed *Drosophila* stocks carrying a *RNAi* transgene and the corresponding *DEP* suppressor mutant on separate autosomes. These stocks were crossed to the *w; GMR-Gal4; UAS-p53* homozygous „tester” stock. Among the F1 offspring, we evaluated the *r.e.* phenotype of the flies which carried the *DEP* suppressor mutant together with the specific *RNAi* silencing construct and the *UAS-p53* transgene, all of them driven by the *GMR-Gal4* driver: *GMR-Gal4>suppr^DEP^*, *UAS-p53*, *UAS-RNAi*.

### RNA preparation and Quantitative Real-Time PCR (RT-qPCR)

Total RNA from twenty heads of 3 day-old *Drosophila* adults for each genetic combination was purified using the RNA isolation kit of Macherey-Nagel (Macherey-Nagel, Düren, Germany) according to the manufacturer’s instructions.1 μg of total RNA was reverse transcribed using the High-Capacity cDNA Archive Kit (Thermo Fisher Scientific, Waltham, MA, USA) according to the manufacturer’s instructions in 20 μl final volume at 37 °C for 2 hours following a pre-incubation at room temperature for 10 min. After inactivating the enzyme at 75 °C for 10 min., the reaction mixture was diluted 30 times. One μl of the diluted reaction mix was used as template in the qPCR. The reaction was performed with gene-specific primers and HOT FIREPol EvaGreen qPCR Mix Plus (ROX) (Solis BioDyne) according to the manufacturer’s instructions at a final primer concentration of 250 nM in Eco Real-Time PCR System (Illumina) under the following conditions: 15 min. at 95°C, 40 cycles of 95°C for 15 sec., 60°C for 20 sec. and 72°C for 20 sec.. Melt curve analysis was done after each reaction to check the quality of the products. Primers were designed online using the Roche Universal Probe Library Assay Design Center or the Integrated DNA Technologies qPCR Assay Design RealTime PCR Tool. Individual threshold cycle (Ct) values were normalized to Ct values of *DmRap*, internal control gene. Relative gene expression levels between induced and control genotypes are presented as fold change values calculated using the formula (fold change=2^ΔΔCt^), according to the ΔΔCt method (Livak and Schmittgen 2001). Primers used in qPCR analysis are listed in **Supplementary Table 2.**

### Statistical analysis of RT-qPCR data

RNA samples were prepared and tested in three parallels (n=3) for each genetic combination. Statistical comparison of normalized Ct (ΔCt) values of control and induced genotypes were done by Student’s t-test (two-tailed, unequal variance). Results are summarized in **Supplementary Table 3**.

## Results

### Isolation and characterization of the mutants carrying the DEP insertions

Overexpression of *Dmp53* in the whole body is lethal. To isolate dominant suppressor mutants of the *p53*-induced apoptosis we took advantage of the *GMR-Gal4* driver, which expresses the Gal4 mainly in the eye (Freeman 1996; Neufeld *et al*. 1998; Ray and Lakhotia 2015). In heterozygous *GMR-Gal4>UAS-p53* flies (*GMR-Gal4/+;UAS-p53/+*) the elevated expression of *p53* causes extensive cell death in the eye imaginal discs, and results in smaller than normal adult eyes with highly disorganized ommatidial arrays; „rough eye”, *r.e.* phenotype, **Fig. 3a** and b; (Ollmann *et al*. 2000; Jin *et al*. 2000). As the flies showing the *r.e.* phenotype are viable and fertile (Kramer and Staveley 2003), we built our activating mutagenesis screen on this approach (for the details, see **Supplementary Fig. 1**). For the mutagenesis, we used the *DEP* P-element construct with two outward-directed *UAS*-coupled promoters („*UAS* promoters”), one at each end (**Fig. 1**). As the Gal4 activates both *UAS* promoters, the transcription simultaneously starts in both directions from the insertion site. The P element preferentially inserts into DNAse hypersensitive regions within or near to gene promoters (Shilova *et al*. 2006). Therefore, we expected that most *DEP* insertions would have been near to the 5’ end of the gene so that the induced downstream transcription would have resulted in enhanced gene expression. We searched for gene mutants (*suppr^DEP^*) which could suppress the *p53* overexpression-induced *r.e.* phenotype when activated by Gal4 in the genetic combination *GMR-Gal4>suppr^DEP^*, *UAS-p53*. Out of more than 2000 insertions on the 2nd and 3rd chromosomes, we recovered seven such mutants (**Figs. 2** and **3**). By sequencing the chromosomal DNA flanking the insertions in the mutant lines, we identified seven genes carrying a *DEP* insertion or associated with it: *Orct2*, *Polr2M, PKA-R2*, *Rga*, *stg/CDC25*, *crol*, *Fak.* In all of them, the *DEP* transposon is inserted near to the 5’ end of the gene (**Fig. 2**): in 3 out of 7, the *DEP* insertions are in the first exon (*Polr2M ^DEP105^, stg^DEP871^, Fak^DEP2107^*). In *Orct2^DEP54^* the *DEP* insert is 129 bp downstream from the transcription start site in the unsplit gene. *Rga^DEP375^* has the insert in the first intron while *crol^DEP1004^* has it in the second exon. In *Pka-R2^DEP327^* the *DEP* transposon is inserted upstream but near to the 5’ end of the gene. As the *DEP* insertion sites are upstream relative to the translation start sites in all but one (*Polr2M ^DEP105^*, 27 bp downstream from the translation start site) of the mutants, we supposed at first that the suppressor effect was a result of the Gal4-induced downstream transcription and overexpression of the gene. However, the transcription starting from a *UAS*-promoter could also spread over to the nearby genes. This assumption was tested by measuring the expression level of the neighbor genes by quantitative PCR and the selective deletion of the *UAS* promoters of the *DEP* element (see below).

**Fig. 1:**
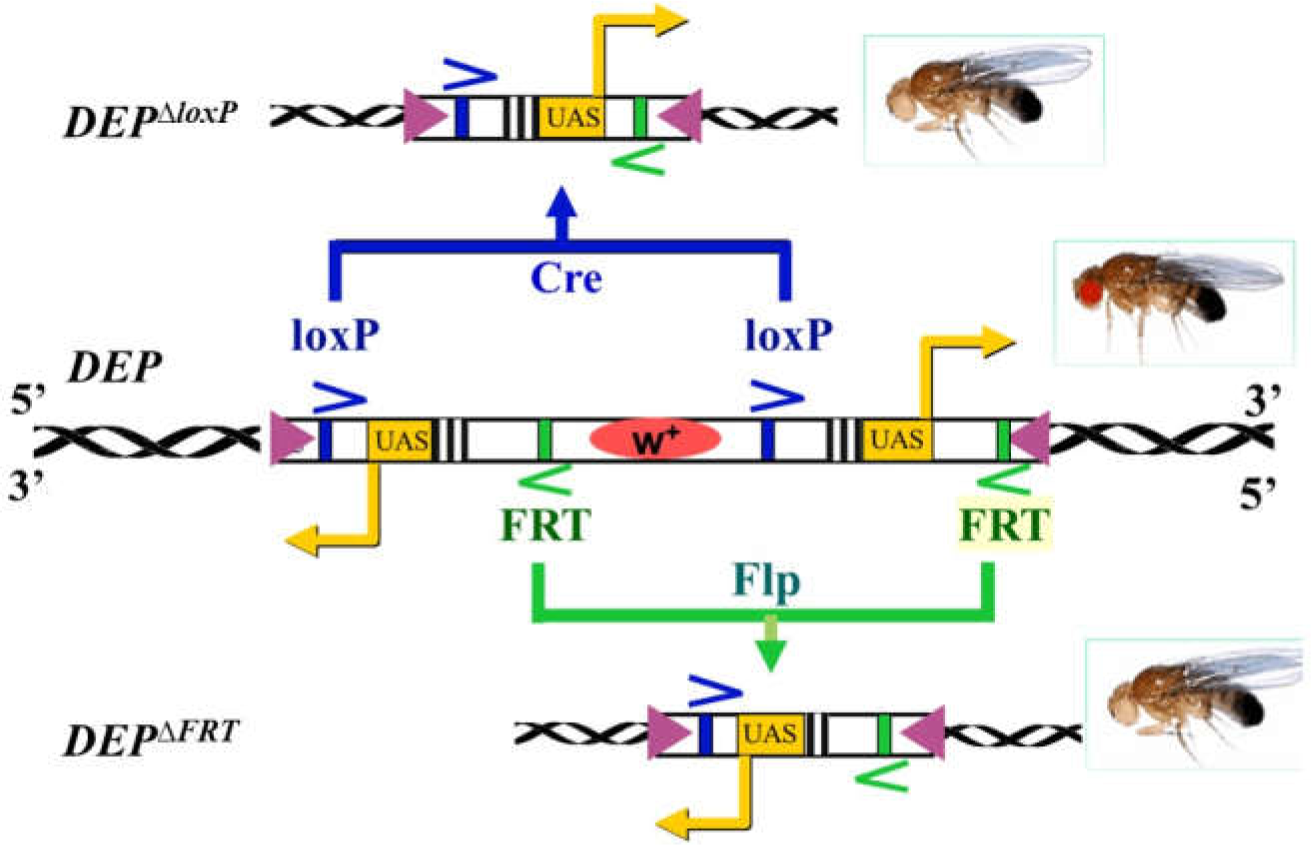
Structure of the *DEP* element and selective deletion of the *UAS* promoters. The two outward-directed *UAS* promoters are located at the ends of the *mini-w^+^ DEP* construct. The *UAS* promoters located at the 5’- and the 3’-ends are flanked by a pair of *FRT* (green) and *loxP* (blue) sites, respectively. Each one of the *UAS* promoters together with the *mini-w^+^*marker can selectively be deleted *in vivo* by the Cre and Flp recombinases (leaving the other *UAS* promoter intact) resulting *ΔloxP* and *ΔFRT* derivatives, respectively. The arrows (yellow) show the directions of the Gal4-induced transcription from the *UAS*-promoters, and the magenta triangles represent the terminal repeats of the *DEP* element.

**Fig. 2:**
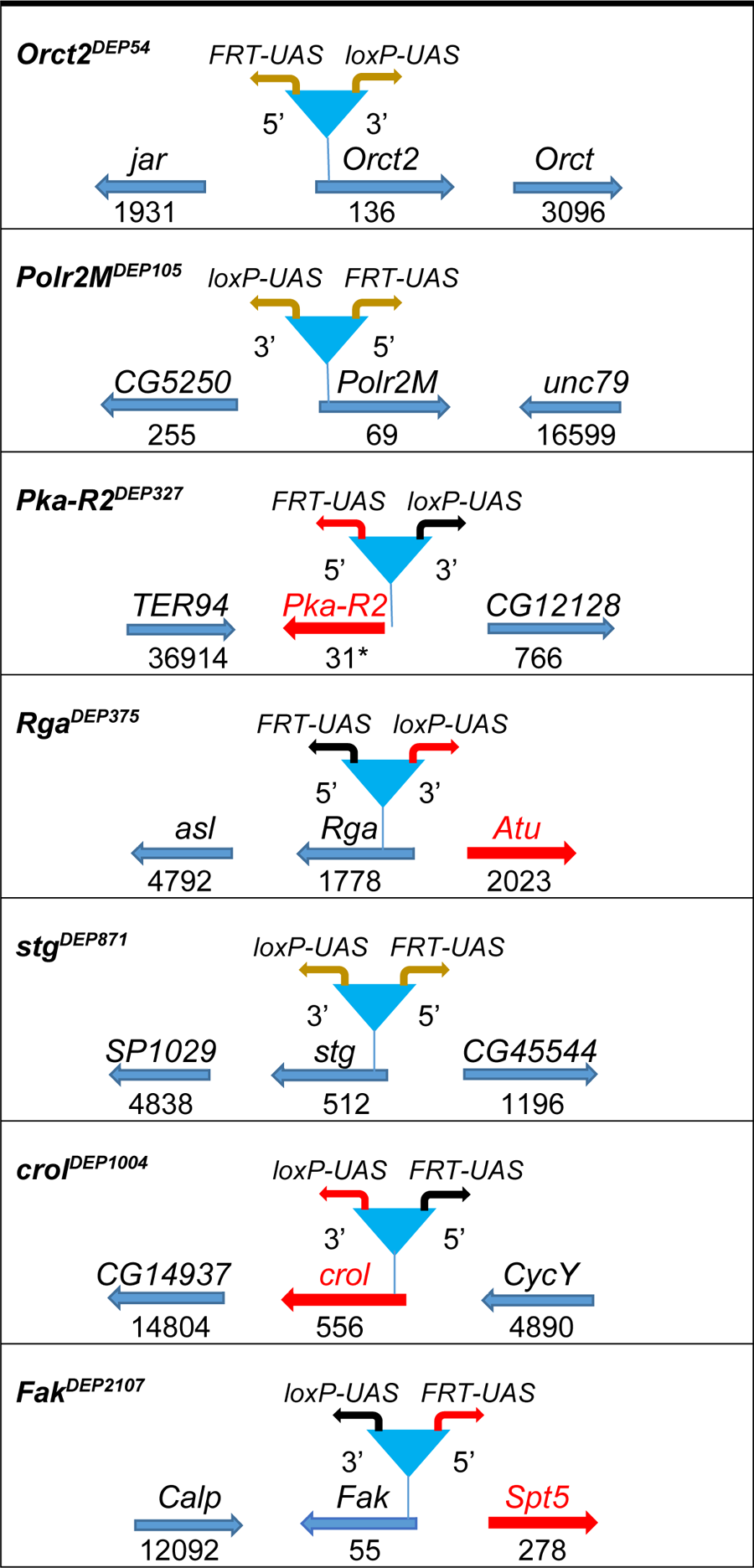
General features of DEP insertions and their genomic neighborhood. For the DNA sequence of the insertion site see Table S1. Arrows label the direction of gene transcription. Numbers below the arrows indicate the distance of the *DEP* insertion site in base pairs downstream from the gene’s transcription start site. Asterisk denotes the distance is upstream from the gene’s transcription start site. Genes in red are responsible for the suppression effect. The blue triangles represent the position of the *DEP* insertions. 5՚*FRT-UAS* and 3՚*loxP-UA*S with red arrows mean the Gal4-activatable *UAS*-promoter identified as the activator of the suppression of apoptosis. Braun arrows mean that the apoptosis suppressor effect of neither *UAS* promoter can be determined unequivocally.

It has to be noted that in all of the different genetic combinations tested we used every component in heterozygous condition. The *GMR-Gal4>UAS-p53* flies showed a strong, characteristic rough eye phenotype which, at the same time, was sensitive enough to be readily modified by the Gal4-activated *DEP* suppressor mutants in the *GMR-Gal4>suppr^DEP^*, *UAS-p53* heterozygous flies (*w^1118^; GMR-Gal4/+; suppr^DEP^*/*UAS-p53*).These heterozygous combinations were able to detect even the weak combined effect of the genes near to the *DEP* insertion (**Fig. 3**).

### Determination of the genes responsible for the apoptosis suppression

As the *DEP* construct carries two outward-directed *UAS* promoters (**Fig. 1**), and the Gal4 simultaneously activates transcription from both of them, we have to define which *UAS* is responsible for the suppressor phenotype. As a first assumption, one would expect that the *UAS* promoter, which initiates downstream transcription of the *DEP*-bearing gene, is responsible for the suppressor effect. Therefore, based on the sequencing data, we determined the orientation of the *DEP* insert, whether the *FRT*-or the *loxP*-flanked *UAS* promoter could initiate downstream transcription of the gene bearing the *DEP* insert. Then the promoters were *in vivo* deleted separately by the FLP or Cre recombinases (see Materials and Methods), and the mutant bearing the truncated *DEP* element (*DEP^ΔFRT^* or *DEP^ΔloxP^*) was crossed to homozygous *GMR-Gal4;UAS-p53* tester flies to see if the apoptosis suppression effect was lost or retained. As the results show, the mutants can be distributed into two groups. In mutants of the first one, if deleting one *UAS* promoter abolishes the suppressor activity, then deleting the other one has weak or no effect. This is true for four mutants (*Pka-R2^DEP327^*, *Rga^DEP375^*, *crol^DEP1004^*and *Fak^DEP2017^*; **Figs. 2** and **3C**). In *Pka-R2^DEP327^*, the *FRT*-deletable *UAS^FRT^* initiating downstream transcription of the gene is responsible for the apoptosis suppression. In the case of *crol^DEP1004^*, the Cre-deletable *UAS^loxP^* promoter initiates the downstream transcription of the *crol* gene and exerts apoptosis suppression. In the case of *Rga^DEP375^* and *Fak^DEP2017^*, the orientation of the *UAS* responsible for the suppression effect points to the upstream direction from the *DEP* insertion, toward a neighbor gene (**Fig. 2**).

In the mutants of the second group, *Orct2^DEP54^*, *Polr2M^DEP105^* and *stg^DEP871^*, deletion of either *UAS* resulted in some sort of a *r.e.* phenotype (**Fig. 3D**). In these cases, we could not assign the suppressor effect unequivocally to one gene or direction, as if it were an additive effect together with the neighbor genes. It has to be noted that, even in the cases when the suppression of apoptosis could be assigned to one gene, the flies having a truncated *DEP* with the „suppressor *UAS”* only showed a weaker suppression of the *r.e.* phenotype than the original mutant bearing the intact *DEP* element (**Fig. 3C**). In these cases the induction of transcription and the possible activation of the neighbor genes was obviously lopsided. This may hint at the possibility that the weaker suppressor effect would either be a result of the missing activity of the neighbor genes on the „silent” side of the truncated *DEP* insert or, if both *UAS*-promoters are simultaneously activated in the intact *DEP* element, there is synergy between them, e.g., by mutually loosening up the chromatin structure, which would enhance the level of transcription of the „suppressor” gene. For the further verification of the effective genes, we used *RNAi* knockdown.

### RNAi knockdown of the effective genes alleviates the apoptosis suppression

**Fig. 3:**
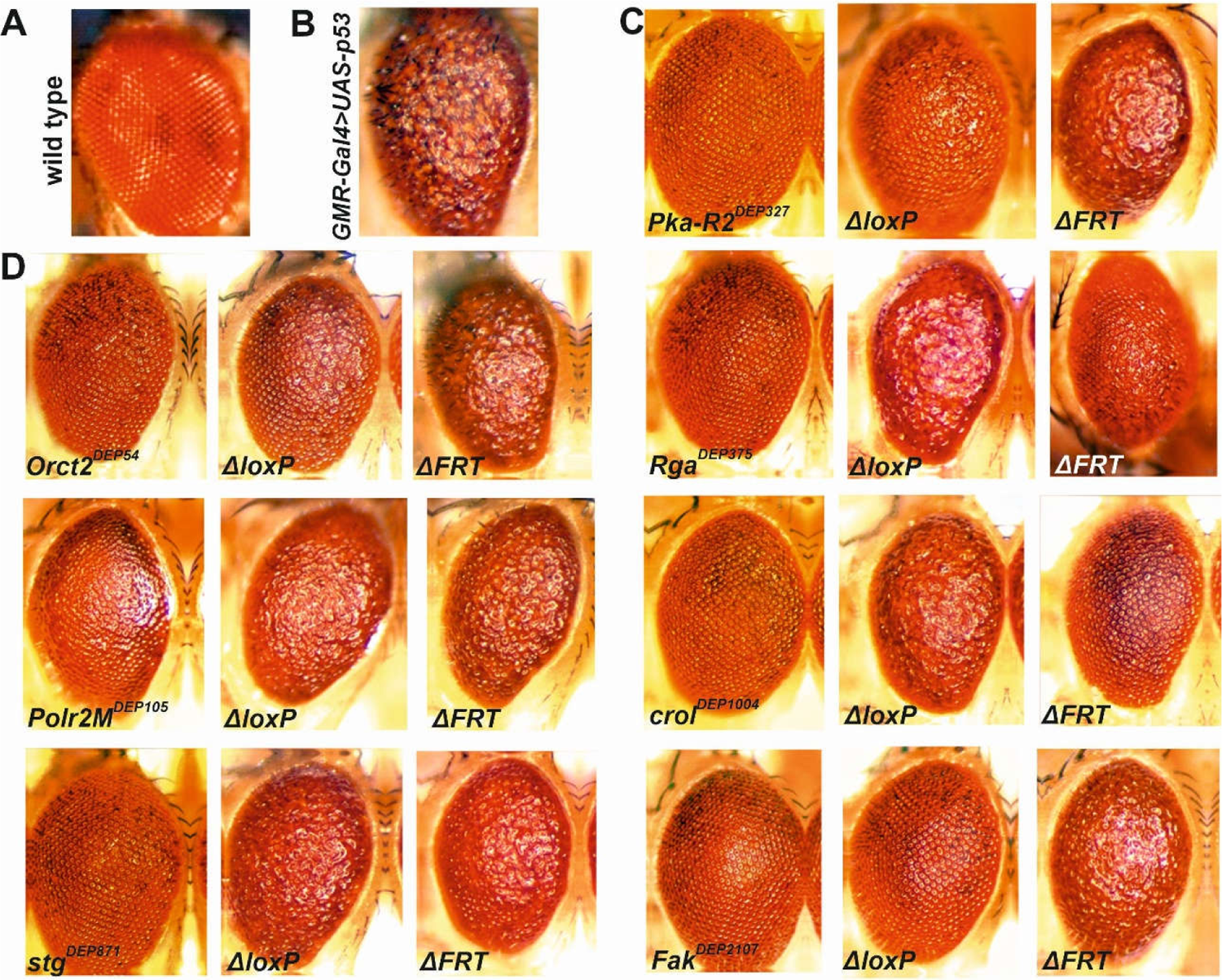
Effect of *Gal4*-activated suppressor gene mutants and their *UAS*-deleted derivatives on the *p53*-induced apoptotic rough eye phenotype. (A) Wild type adult eye. **(B)** Rough-eye phenotype of the *GMR-Gal4>UAS-p53*. **(C**-**D)** Apoptosis suppression effect of *GMR-Gal4>suppr^DEP^, UAS-p53* combinations. *ΔloxP* and *ΔFRT* stand for the *UAS*-deleted *DEP* mutant derivatives *DEP^ΔloxP^* and *DEP^ΔFRT^,* respectively (see Figure 1). **(C)** Suppression of rough eye phenotype is caused by one of the two *UAS*s. **(D)** Deletion of either *UAS*-promoter results in the same rough-phenotype, i.e., the apoptosis suppression effect cannot be definitely related to either *UAS*-promoter.

In the *GMR>suppr^DEP^+UAS-p53* adult flies the apoptotic rough eye phenotype is suppressed by the activated *DEP* insertion. However, if the fly additionally carries an *UAS-RNAi* transgene which specifically silences the suppressor gene e.g., *GMR-Gal4>UAS-RNAi, suppr^DEP^, UAS-p53,* i.e.*, w^1118^; GMR-Gal4/UAS-RNAi; suppr^DEP^/UAS-p53*, the rough eye phenotype can be partly or entirely restored (**Table 1**). To this end, a series of stocks were established carrying a *DEP* mutant and a *UAS-RNAi* transgene which specifically silences the *DEP*-bearing gene or one of the neighbor genes. After crossing these stocks to the *GMR-Gal4; UAS-p53* tester combination, we evaluated the eye phenotype of the F1 flies which carried together the *DEP* suppressor mutant, its specific *RNAi* silencing transgene and the tester combination. Gal4-induced expression of the *RNAi* transgenes in themselves did not cause *r.e.* phenotype (not shown). The results are summarized in **Table 1**.

**Table 1:** Effect of RNAi silencing the apoptosis suppressor effect in the *GMR>suppr^DEP^+UAS-RNAi+UAS-p53* combination. ^1^Only those RNAi constructs were tested which were located on different chromosomes from the *DEP* insertions. Specification of the RNAi stocks is given with the stock center name and stock number. The numbers in brackets mean the score of the rough-eye phenotype. (see **Supplementary Fig. 2**) Abbreviations: BDSC: Bloomington Drosophila Stock Ctr. (Bloomington, Indiana); VDRC: Vienna Drosophila Resource Ctr. (Vienna, Austria); NIG-FLY: National Institute of Genetics (Mishima, Japan).

| **DEP insertion mutant** | **Tested genes in the DEP insertion region** | **Effective RNAi constructs<sup>1</sup> (Score>0)** | **Ineffective RNAi constructs<sup>1</sup> (Score=0)** |
| --- | --- | --- | --- |
| *Orct2*<sup>DEP54</sup> | *Orct* |  | VDRC 106681<br>BDSC 57583 |
|  | *jar* |  | VDRC 37534<br>VDRC 37535<br>VDRC 108221<br>BDSC 28064 |
| *Polr2M*<sup>DEP105</sup> | *PolR2M* | BDSC 42917 (1-2) |  |
|  | *CG5250* | BDSC 57432 (1-2) |  |
| *Pka-R2*<sup>DEP327</sup> | *Pka-R2* | NIG-FLY 15862R2 (4)<br>BDSC 27680 (3-4)<br>BDSC 34983 (4) |  |
|  | *TER94* |  | BDSC 31968 |
|  | *CG12128* |  | BDSC 33997 |
|  | *CG1407* |  | BDSC 50601 |
| *Rga*<sup>DEP375</sup> | *Rga* |  | BDSC 57549 |
|  | *asl* |  | BDSC. 38220 |
|  | *Atu* | VDRC 106074 (2-3) |  |
|  | *Spec2* |  | BDSC 65206 |
| *stg*<sup>DEP871</sup> | *stg* | BDSC 36094 (2) |  |
| *crol*<sup>DEP1004</sup> | *crol* |  | BDSC 44643 |
|  | *CG14937* |  | BDSC 31483 |
|  | *CycY* |  | BDSC 34009 |
|  | *esc* |  | BDSC 31618 |
| *Fak*<sup>DEP2107</sup> | *Fak* |  | VDRC 17957<br>BDSC 29323<br>BDSC 33617<br>BDSC 35357 |
|  | *CalpA* |  | BDSC 29455 |
|  | *Spt5* | NIG-FLY 7626R-3 (3) |  |
|  |  | BDSC 34837 (4) |  |

In the case of *Pka-R2^DEP327^* the results were straightforward: we tried three different *RNAi* transgenes silencing *Pka-R2*, and all of them restored the *r.e.* phenotype verifying that the suppressor effect was really caused by the overexpression of *Pka-R2.* At the same time, silencing the neighbor genes *TER 94*, *CG12128* and *CG1407* had no effect (**Table 1**). In the case of *Fak^DEP2107^*, the *DEP* element is inserted in the *Fak* (*Focal adhesion kinase*) gene, and the *loxP*-deletable *UAS* activates its ectopic expression (**Fig. 2**). It has to be noted that the mammalian ortholog FAK is a potent apoptosis suppressor (Sonoda *et al*. 2000; Kurenova *et al*. 2004). However, deletion of the *UAS^loxP^* did not restore the apoptosis suppressor effect (Figure 3), and the *Gal4*-induced expression of a *UAS-Fak* transgene remained ineffective either (not shown). Instead, deleting the upstream-oriented *UAS^FRT^* at the 5^’^end of *DEP* abrogated the *r.e.* suppression suggesting that the activity of the *UAS^FRT^*promoter was responsible for the suppressor effect. The *UAS^FRT^*in this case points in the 5’ direction to the nearby *Spt5* gene indicating that, in agreement with the promoter deletion test, this gene can be responsible for the apoptosis suppression. In addition, we tried a *UAS-Spt5* transgene which effectively suppressed the *r.e.* in the *GMR-Gal4*>*UAS-Spt5, UAS-p53* combination (not shown).

In addition, *RNAi* knockdown of the *Spt5* expression by *UAS-RNAi* transgenes in the *GMR-Gal4*>*Fak^DEP2107^+UAS-Spt5i+UAS-p53* combination brought back the rough eye (**Table 1**). All these results prove that the *Spt5* gene is an apoptosis-suppressor.

In the case of *Rga^DEP375^*, the *UAS^loxP^*pointing in the direction of the *Atu* gene shows the suppressor activity (**Fig. 2**). In accordance with this, the *Atu*-silencing *RNAi* transgene restored the *r.e.* phenotype but silencing *Rga* and the neighboring genes *asl* and *Spec2* had no effect (**Table 1**). In *stg^DEP871^*, the *DEP* element sits in the first exon near to the 5’ end of *stg*, and the deletion test showed that, to some extent, both *UAS* promoters were responsible for the apoptosis suppression. The *UAS^FRT^* initiates transcription toward *CG45544*, an unknown gene nearby (**Fig. 2**). There was no *RNAi* construct available for this gene so we could not test the possible influence of *CG45544* on the suppressor effect. On the other hand, *stg* is the *Drosophila* ortholog of the *cdc25* phosphatase, a key factor of mitosis progression, and previous studies showed that *cdc25* was able to inhibit apoptosis (Kylsten and Saint 1997; Fuhrmann *et al*. 2001; Cho *et al*. 2015). Therefore, we tested the *UAS-stg* transgene to see whether it could suppress the *r.e.* phenotype in the *UAS-stg/GMR-Gal4; UAS-p53/+* combination. The test showed that the expression of *Drosophila stg* was able, albeit weakly, to inhibit, the *p53*-induced apoptosis (not shown). Taken together, one can suppose that in *stg^DEP871^* the simultaneously induced expression of *stg* and *CG45544* could additively suppress apoptosis.

In the case of *Polr2M^DEP105^*, the *DEP*-bearing *Polr2M* and the neighbor gene *CG5250* were separately silenced. As it revealed, both tested *RNAi* transgenes weakened the suppression of apoptosis to some extent, but their effect was not strong (Table 1). This suggests an additive suppressive effect of the two genes.

### A RT-qPCR survey of gene activation by the GMR-Gal4 driver

Supposing that the Gal4-induced overexpression of the gene bearing the *DEP* insertion can spread over the neighbor genes in the region, and their elevated expression may also contribute to the suppressor phenotype, in a RT-qPCR experiment we systematically tested the expression levels of the nearby genes as well. The results of this survey are summarized in **Fig. 4, Supplementary Fig. 3** and **Supplementary Table 3.**

In addition to each gene with the *DEP* insert, Table S3 contains the genes in the surrounding region, and shows the distances in kbp between the genes’ transcription start sites and the *DEP* insertion site, as well as the fold change values of their *GMR-Gal4*-induced expression levels. The expression levels were measured with *UAS-p53* in the background (*GMR-Gal4>suppr^DEP^, UAS-p53* versus *GMR-Gal4>UAS-p53*). For the statistical evaluation of the results see **Supplementary Table 3**.

The *DEP* as a *P* element derivative, mostly inserts itself into or near to the 5’ end of the genes. Consequently, one of the two *UAS*-promoters can always start the downstream transcription of the gene resulting in a supposedly normal mRNA. Compared to the uninduced „basic activity”, *GMR-Gal4* can induce a significantly elevated expression of the *DEP*-bearing gene, and the activating effect can spread to the nearby genes as well. In general, the level of activation decreased with the growing distance from the *DEP* insert but the actual values varied depending on the gene and the region (**Fig 4, Supplementary Fig. 3** and **Supplementary Table 3**).

The fold degree of activation largely depended on the basic, uninduced level of the gene activity (see in FlyAtlas, www.flyatlas2.org): when it was very low in general, the Gal4-induced expression could reach high or extremely high relative levels. *E.g*. for *stg^DEP87^*^1^ the *GMR-Gal4*-induced fold change was 146x while in the vicinity, *CG45544* (distance from *DEP* insertion site 1.6 kb) and *CR45568* (distance 21.8 kb) were induced by 7x10^4^ and 4x10^3^ times, respectively (**Fig. 4, Supplementary Fig. 3 and Supplementary Table 3**).

**Fig. 4:**
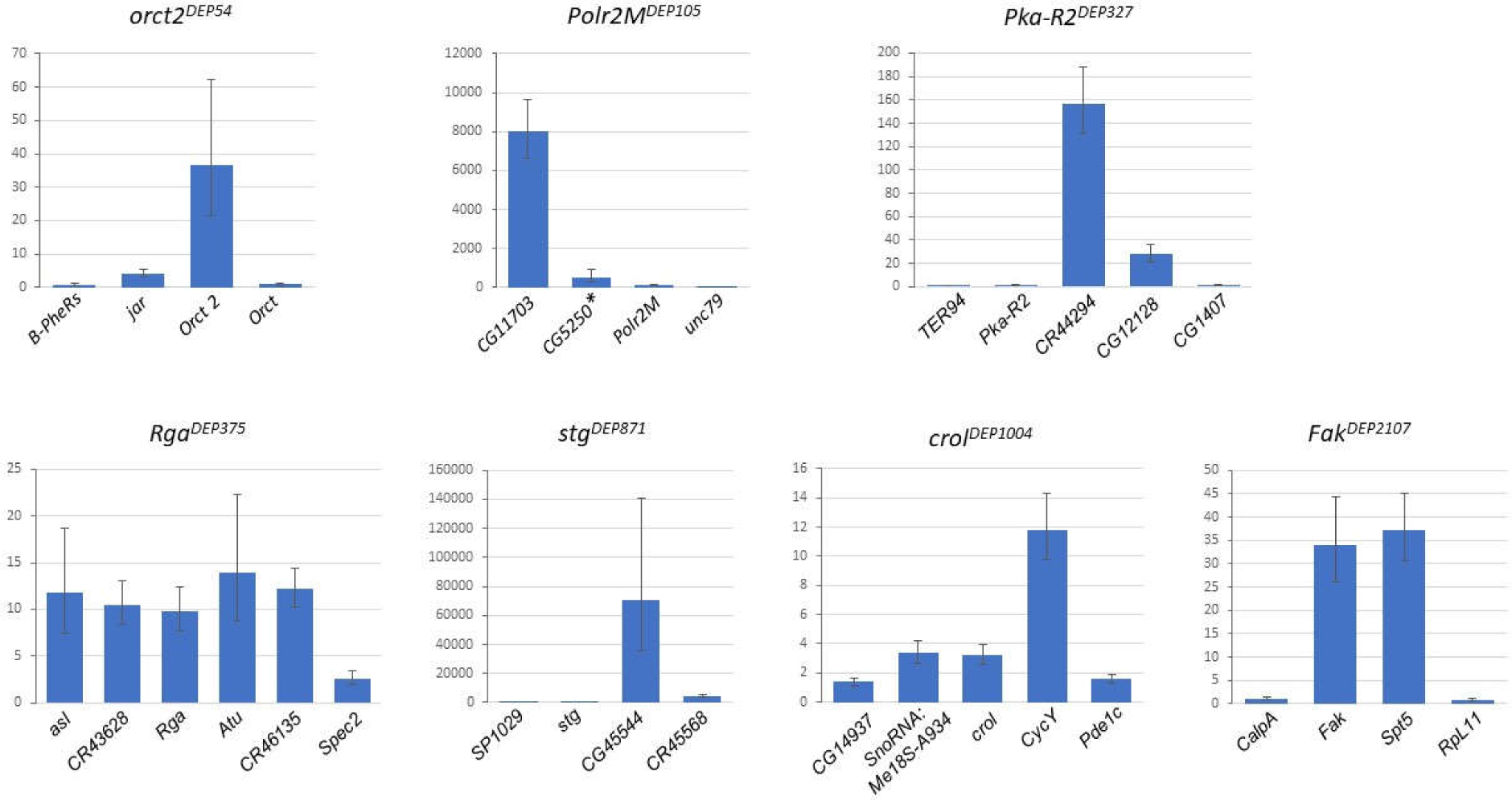
Gal4 induced activity of the genes bearing the *DEP* insertion and the genes in the neighborhood. The columns represent the fold change of the gene expression measured in *GMR-Gal4*>*Suppr^DEP^, UAS-p53* versus *GMR-Gal4>UAS-p53*. For the numerical results see Table S3. *: Uninduced control: *w; Polr2M^DEP105^/TM3*.

It is worth to note that, in general without Gal4 induction, the presence of the *DEP* insert alone did not increase the expression level of the *DEP*-bearing gene and its neighbors in the region. There are a few exceptions, however: in the vicinity of the uninduced *Polr2M ^DEP105^*, the *CG11703* gene (distance from *DEP* insertion site 1.3 kb) showed a 20x increase in comparison to its basic expression level in homozygous *w* with no *DEP* insertion. In *stg^DEP871^*, the uninduced basic expression level of *stg* was 2x higher than that without the *DEP* insertion. Similarly, the nearby *CG45544* (distance from *DEP* 1.6 kb) and *CR45568* (distance 21.8 kb) genes showed almost 40x and 9x increased expression, respectively. As mentioned above, these genes show very low basic expression in the adult head and brain so even a small increase in their expression can relatively be highly magnified in the fold change. Finally, in *Fak^DEP2107^*, the insert-bearing *Fak* gene has a 4x increased expression without Gal4 induction.

Theoretically, if the Gal4-induced transcription starting from the *UAS*-promoter runs opposite to a gene’s own transcription, it could interfere with the production of mRNA, e.g. through an *RNAi*-type of interaction between the two transcripts. As **Supplementary Table 3** shows, we could not observe any obvious example of such an effect.

### Apoptosis suppressor mutants in co-expressed gene clusters

In the genomes of eukaryotic organisms there are groups of neighbor genes which are co-expressed. In the genome of *Drosophila melanogaster* Spellman and Rubin identified more than 200 such gene clusters (Spellman and Rubin 2002). It seems to be a logical assumption that in these neighborhoods the co-expressed genes with completely different functions cooperate for establishing some complex cellular entities like a complex of different proteins, a biochemical pathway, an elaborate cellular function, etc.

In our mutants, we compared the chromosomal location of the genes in the *DEP* insertion neighborhoods with that of the known co-expressed gene clusters in the *Drosophila* genome (Spellman and Rubin 2002). It revealed that in three out of seven mutants (*Polr2M^DEP105^*, *Rga^DEP375^*and *Fak^DEP2107^*), the groups of the *DEP-*bearing genes and their neighbors were included in three separate co-expressed clusters (**Supplementary Table 4**). It should mean that even without Gal4 activation, the genes in these *DEP* insertion neighborhoods are co-expressed with the other genes of the cluster. In addition, two other mutants, *Orct2^DEP54^*and *crol^DEP1004^* are located near to the boundary of separate co-expressed clusters (**Supplementary Table 4**). With the *GMR-Gal4* driver we induce the expression of the genes in the *DEP* insertion neighborhoods only. In the case of *Polr2M^DEP105^*the Gal4-induced co-expression revealed the anti-apoptotic ability of the genes in the near neighborhood of the *DEP* insertion. Whether the other genes in the cluster have similar ability or could influence the anti-apoptotic activity of the *DEP* neighborhood genes remains to be seen.

## Discussion

Apoptosis is the primary defense mechanism against cancer, and *p53*, the chief regulator of the apoptotic process, is one of the most important tumor suppressor genes. *p53* is a transcription factor which, directly or indirectly, regulates the expression of hundreds of genes, and is itself regulated by upstream genetic factors. Negative regulation of apoptosis and/or *p53* plays important roles in promoting tumor development. Large-scale screens in mammalian cell lines identified numerous genes potentially inhibiting apoptosis but for many of them their significance and mechanism of action are still not clear (Kirsch and Kastan 1998; Vousden and Prives 2009; Portt *et al*. 2011; D’Orazi 2021).

In the present study we used the advanced genetic system of *Drosophila melanogaster* to identify and recover new apoptosis suppressor genes. We made an activation screen by using the novel *DEP P*-element construct which can initiate Gal4-inducible transcription to both directions from the insertion site and, by selectively deleting *in vivo* one or the other *UAS*-regulated promoter, one can determine the direction from which the apoptosis-suppressor effect originates. In addition to the gene bearing the *DEP* insertion, one can also test the effect of the neighbor genes in this way. In the *GMR-Gal4* (Glass Multimer Reporter) driver, the Gal4 is mainly expressed in the developing eye disc and the adult eye (Roman and Davis 2002; Yang *et al*. 2005). However, *GMR-Gal4* expression was detected in other tissues as well, namely in brain, trachea, and leg discs (Li *et al*. 2012). In addition, Ray and Lakhotya (2015) found that the strong *Gal4* expression on its own can interfere with normal eye development resulting in some „rough eye” adult phenotype. To avoid these possible disturbing effects, we used only one copy of the *GMR-Gal4* driver in heterozygous condition which on its own did not interfere with the normal eye development in the genetic combinations used. However, if we have only one copy of *GMR-Gal4* in the combination, the number of the Gal4 binding sites can become critical. If too many *UAS* motifs compete for the limited amount of Gal4 protein, one could expect that the Gal4-induced apoptosis and *r.e.* phenotype would become weaker mimicking the suppression of apoptosis. In the „tester” combination *GMR-Gal4>suppr^DEP^, UAS-p53* where the single *GMR-Gal4* drives a *DEP* inserted element and the *UAS-p53* together, there are three *UAS* motifs sharing the Gal4 and show the usual *r.e.* phenotype (Figure 2). If the combination contains four *UAS*s (e.g., *GMR-Gal4>suppr^DEP^, UAS-RNAi, UAS-p53)*, the *r.e.* is still well visible (**Table 1**). Above this number, however, the *r.e.* phenotype begins to weaken. Hence, any of the genetic combinations, where the *GMR-Gal4* driver was used, contained only four *UAS*s at the maximum.

When the activation of a *DEP* suppressor mutant by the *GMR-Gal4* results in the suppression of the apoptotic rough eye phenotype in the *GMR-Gal4>suppr^DEP^, UAS-p53* combination, the easiest explanation would be that the suppressor effect is resulted by the elevated expression of the gene bearing the *DEP* insert. As the P element and its derivatives tend to insert at the 5’ end of the gene, the downstream-directed *UAS*-promoter of the *DEP* insert would induce the gene expression resulting in the apoptosis suppression. In such a case, deletion of the downstream-directed *UAS*-promoter should eliminate the suppressor effect and bring back the rough eye phenotype. This is the case e.g., for *Pka-R2^DEP327^* (**Figs. 2** and **3**). This is further supported by the fact that the *RNAi* constructs specifically decreasing the activity of *Pka-R2* also eliminate the apoptosis suppression in the tester combination *GMR-Gal4>Pka-R2^DEP327^, UAS-RNAi, UAS-p53*, and restore the rough eye (**Table 1**). In the case of *crol^DEP1004^*, the situation is similar: as the selective deletion of the *DEP UAS*-promoters shows, the direction/orientation of the „suppressor” *UAS*-promoter in the *DEP* insert matches the downstream direction of the *crol* gene (**Figs. 2** and **3**) although the suppressor effect of *crol^DEP1004^* was not eliminated by the *crol*-specific *RNAi* construct we tested (**Table 1**).

In two other mutants (*Rga^DEP375^* and *Fak^DEP2107^*) the *UAS*-promoter responsible for the suppressor effect was directed towards the *Atu* and *Spt5* genes in the neighborhood, respectively, suggesting that these genes may be responsible for the apoptosis suppression (**Figs. 2** and **3**). This conclusion was further supported by the *RNAi* results: in both cases the *p53*-induced apoptosis was suppressed by the *Atu*- and *Spt5*-specific *RNAi* constructs (**Table 1**). In addition, the apoptosis suppression could also be achieved by the *GMR-Gal4*-induced expression of a *UAS-Spt5* construct (not shown).

For the remaining three mutants *orct2^DEP54^, Polr2M^DEP105^*and *stg^DEP^*^871^, however, the *UAS*-deletion gave equivocal results: deleting one or the other *UAS* did not abolish unequivocally the apoptosis suppression effect, significant *r.e.* phenotype appeared in either deletion variant (**Figs. 2** and **3**). The *RNAi* gene silencing gave similar equivocal results for *Polr2M^DEP105^*. Silencing the *Polr2M^DEP105^* gene or its close neighbor *CG5250* restored a weak apoptotic rough eye phenotype in both cases (**Table 1**).

In the case of *stg^DEP871^*, deleting either *DEP* promoter showed the same result: a weak suppression of the rough eye phenotype (**Fig. 3**). The *RNAi* silencing of *stg^DEP871^* also resulted in a moderate apoptotic eye phenotype (**Table 1**). As no matching *RNAi* construct was available on the 2nd chromosome for the neighbor gene *CG45554,* we could not test the effect of its silencing. Ruiz-Losada and her co-workers (Ruiz-Losada *et al*. 2022) recently reported on an observation which seemingly opposes our above conclusion: they found that over-expressing the *stg* gene in larval wing discs for 24 hrs followed by X-ray irradiation (IR) significantly increased the number of apoptotic cells compared to IR with no previous *stg* induction. They concluded that *stg* is a pro-apoptotic gene. Furthermore, they studied the connection of the Cyclin-dependent kinase 1 (Cdk1) and p53 proteins in the G2/M transition of the cell cycle, and found that the two can make a complex, and Cdk1 promotes the binding of p53 as a transcription factor to the specific binding sites. This can suggest an alternative explanation to the role of *string*: during the 24 h over-expression, the stg protein as a tyrosine protein phosphatase, directly dephosphorylates and activates Cdk1 which after the IR exerts a strong contribution to the activation of p53 and apoptosis. The connection of *stg* to apoptosis needs further investigation.

Taken together, in addition to the *DEP* mutant gene, at least two (maybe more) neighbor genes in cooperation seem to be responsible for the suppression of apoptosis in the above cases, and we suppose that the normal or nearly normal eye phenotype is the result of their additive effect. For this effect, however, the expression of these genes has to be induced which is the result of the activating effect of the *DEP* insert spreading over to these neighbor genes. Therefore, we measured with quantitative PCR the rate of expression of the genes around the *GMR-Gal4*-activated *DEP* insertions. **Fig. 4** and **Supplementary Table 3** show the fold change of expression level of the genes in *GMR-Gal4>suppr^DEP^, UAS-p53* compared to the expression in *GMR-Gal4>UAS-p53.* As the data show, the level of transcription is enhanced not only for the gene bearing the *DEP* insert but it spreads over several kilobases to the neighbor genes in the vicinity. In general, the level of transcriptional activation is higher in the genes close to the *DEP* insertion site and decreases with the distance (**Supplementary Fig. 3** and **Supplementary Table 3**).

Extremely high relative activity/fold change can be seen in the activated genes which have very low uninduced basic activity in the adult head. This is the case e.g., for *stg^DEP871^*, where the *stg* gene and its upstream neighbors have very low basic activity in the adult head (FlyAtlas, www.flyatlas2.org) and, accordingly, show high relative activity when induced by *GMR-Gal4* (**Fig. 4** and **Supplementary Table 3**). One may also speculate that a relative stability of the mRNAs of the induced genes in the adult head could also contribute to the Gal4-induced accumulation of specific mRNAs.

The primer sequences we used for the RT-qPCR corresponded to the normal mRNA sequences for all the genes tested including those ones in which the direction of their own transcription was opposite to the orientation of the Gal4-induced *UAS* promoter. This should mean that the data shown in **Fig. 4** and **Supplementary Table 3** represent the Gal4-induced transcription of the neighbor genes started from their own promoters. The cause of the spreading gene activation effect can be the probable loosening up of the local chromatin structure by the parallel transcriptional activity of the *DEP UAS*-promoters. We suppose that while the *GMR-Gal4*-induced direct transcription can not reach the neighbor genes, the loosening up of the chromatin structure can spread further through the region and elevates the accessibility and the expression level of the surrounding genes. This may be the mechanism by which the *GMR-Gal4*-activated *DEP* element indirectly activates the neighboring genes, and, in certain cases, their weak activities are added up in a significant effect suppressing apoptosis. To our knowledge, this is the first observed example of such additivity/cooperativity of neighbor genes in the suppression of the *p53*-induced apoptosis.

On the chromosomes of eucaryotic organisms there are gene clusters in which the genes are co-expressed (Ben-Shahar *et al*. 2007; Michalak 2008; De and Babu 2010; Mihelčić *et al*. 2019). Some of the clusters contain genes with similar functions, while others have genes with diverse functions (Lercher *et al*. 2002). Such co-expressed gene clusters were found also in *Drosophila* (Boutanaev *et al*. 2002; Spellman and Rubin 2002; Stolc *et al*. 2004).

As it revealed, three out of the seven mutants (*Polr2M^DEP105^*, *Rga^DEP375^* and *Fak^DEP2107^*) and their neighbor genes are located in three separate co-expressed clusters, and two mutants (*Orct2^DEP54^*and *crol^DEP1004^*) are outside but very near to other separate clusters. (Table S4). This is particularly interesting in the case of *Polr2M^DEP105^*, in which the apoptosis suppression seems to be an additive effect of the activated genes around the *DEP* insertion (see above). This means that the co-expression is the normal function of these genes, and the control of apoptosis could prevail without the *GMR-Gal4* activation either. Whether the other genes in the cluster would exert similar effect, and the genes in the other clusters above could influence the suppression of apoptosis, needs further investigation.

The question promptly arises whether these genes near to the *DEP* insertion site can really suppress apoptosis or they can influence the regulation of apoptosis in any respect. To this end, we made a survey in the literature for the genes tested in the RT-qPCR experiment. As the *Drosophila* genes are not so well-characterized in this respect, we examined their human orthologs as well. The programmed cell death is one of the most important factors blocking the development of cancer, therefore a lot of information and data can be found about the protein-coding human genes in this respect. Logically, if a gene has any anti-apoptotic effect, its over-expression promotes cell proliferation and tumor development while its reduced activity has the opposite effect. **Supplementary Table 5** shows the 7 *DEP*-bearing *Drosophila* genes and their close neighbors (26 genes) as well as their human orthologs (24 genes). Altogether, according to the literature, twenty-one of these human genes possibly have anti-apoptotic activity, two genes are pro-apoptotic and one gene is uncertain in this respect. Our results in *Drosophila* call the attention to the apoptosis suppressive effect of these genes.

As our results suggest, in certain cases not only the gene examined but other genes in the vicinity can also influence the regulation of the programmed cell death, especially if they are overexpressed ectopically. While the effect of a single gene can be negligible, the combined effect together with the neighbor genes can add up to a significant level. This aspect should also be taken into consideration when studying the genetic regulation of apoptosis or cancer in which more than one gene in a chromosomal region are activated or extenuated together.

## Data availability statement

The strains and the DEP transposon are available upon request. All data confirming the conclusions of the article are included in the article, figures and tables.

## Acknowledgements

This work was supported by the Hungarian Scientific Research Fund (OTKA K69279). BMM and IK were supported by the German Research Foundation (DFG)-Hungarian Academy of Sciences (MTA) Collaboration Program (UNG 436 113/81/0-6). AF was supported by NKFIH 138128. MZ was supported by the European Community’s Program Interreg Bayern Tschechische Republik (BYCZ01-039). We are grateful to Mária Kopp, Tünde Tóth and Rozália Török for technical assistance.

## Author contribution

TL and IK designed and constructed the DEP element. IK and TS directed the experimental work. ÉB, OM, IH, EM, EV and BK isolated the DEP insertion mutants and their UAS-deleted derivatives. IK, ÉB and IB designed and performed the p53-suppressor screen. TL, KS, IT and BMM determined the position of DEP-insertion sites. BR, LMM and ZA performed the RNAi experiments. TS, YHL, ÁZ, EV, LP and MZ designed and performed the RT-qPCR experiments. ZH, TS and AF performed bio-informatic and statistical analysis. EV edited the figures and tables. IK, TS and MZ and wrote the manuscript.

